# Synthetic soil crusts against green-desert transitions: a spatial model

**DOI:** 10.1101/838631

**Authors:** Blai Vidiella, Josep Sardanyés, Ricard V. Solé

## Abstract

Semiarid ecosystems are threatened by global warming due to longer dehydration times and increasing soil degradation. Mounting evidences indicate that, given the current trends, drylands are likely to expand and possibly experience catastrophic shifts from vegetated to desert states. Here we explore a recent suggestion based on the concept of ecosystem terraformation, where a synthetic organism is used to counterbalance some of the nonlinear effects causing the presence of such tipping points. Using an explicit spatial model incorporating facilitation and considering a simplification of states found in semiarid ecosystems i.e., vegetation, fertile and desert soil, we investigate how engineered microorganisms can shape the fate of these ecosystems. Specifically, two different, but complementary, terraformation strategies are proposed: *Cooperation*-based: *C*-terraformation; and *Dispersion*-based: *D*-terraformation. The first strategy involves the use of soil synthetic microorganisms to introduce cooperative loops (facilitation) with the vegetation. The second one involves the introduction of engineered microorganisms improving their dispersal capacity, thus facilitating the transition from desert to fertile soil. We show that small modifications enhancing cooperative loops can effectively change the location of the critical transition found at increasing soil degradation rates, also identifying a stronger protection against soil degradation by using the *D*-terraformation strategy. The same results are found in a mean field model providing insights into the transitions and dynamics tied to these terraformation strategies. The potential consequences and extensions of these models are discussed.

## I. INTRODUCTION

Global warming is changing the dynamics and resilience of ecosystems, damaging many of them and creating the conditions for widespread diversity loss [2, 34, 35]. Because of the presence of nonlinear effects, many ecosystems display so called tipping points associated to community collapse. Among these systems, drylands (which comprise arid, semi-arid and dry-subhumid ecosystems) are a specially fragile subset of major importance: they include more than 40% of terrestrial ecosystems and host a similar percentage of the current human population [31]. Increasing aridity is pushing these ecosystems towards serious declines in microbial diversity, land degradation, loss of multifunctionality as a desert state is approached [23, 31]. Dedicated efforts have been addressing several avenues to both understanding how transitions can be anticipated by means of warning signals [16, 18, 39, 45] and even prevented [31, 40, 46].

Drylands are characterised by the presence of organisms that have adapted to low moisture availability, damaging UV radiation and high temperatures [29]. A rich community structure and the maintenance of physical soil coherence are essential to prevent drylands from degradation. In this context, some universal types of interactions that occur among species in arid habitats inevitably lead to breakpoints associated to the existence of multiple alternative states, identified in field data [4]. A well known class of these interactions takes place among vascular plants and is known as *facilitation* [5, 7, 48], i. e. non-trophic interactions between individuals mediated through changes in the abiotic environment or through other organisms favouring individual growth and reproduction [8, 9, 36].

A common outcome of facilitation is the emergence of spatial patchiness (see Fig. 1(a-b)). Such patterns are often remarkably organised in space [19]. These transitions can be sometimes observed in the same landscape under the presence of environmental gradients [5]. In this context it has been conjectured [17, 18] that spatial correlations can be used as indicators of forthcoming green-desert shifts [16]. The presence of catastrophic transitions deeply modifies our perception of risks associated to climate change and land degradation. Once a critical state is approached, unstoppable runaway processes are unleashed [38]. Despite much progress has been made in modelling drylands [10, 11, 41] as well as in identifying warning signals [16, 18, 39, 41, 45], feasible strategies to prevent green-desert transitions are currently scarce.

**FIG. 1.**
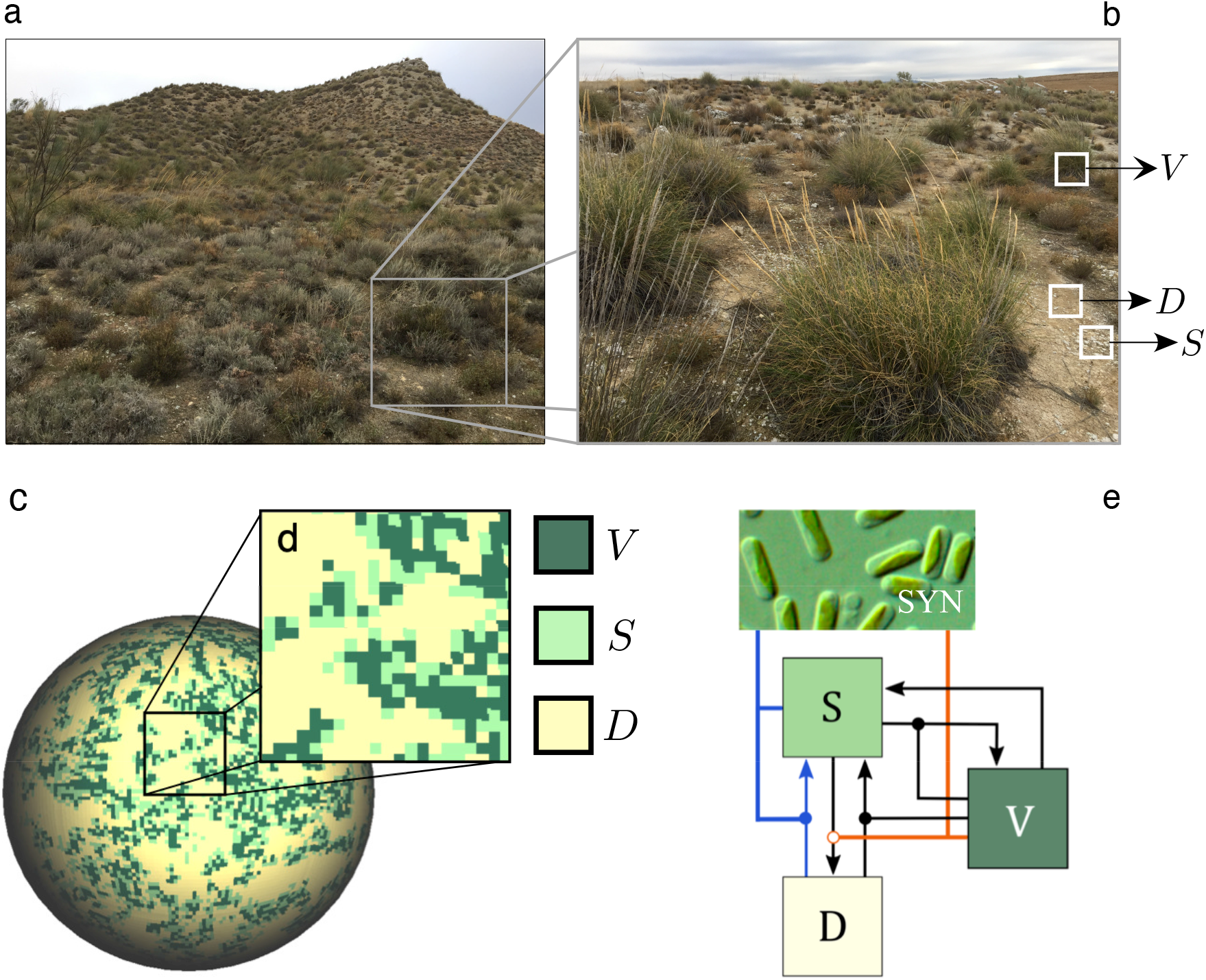
Complexity and local states in semiarid ecosystems. (a) Example of a semiarid landscape from Aranjuez (central Spain). An enlarged view is shown in (b) showing patches of vegetation, fertile soil with crust and bare soil. These three states allow to define three different classes of patches, namely vegetation (*V*), functional soil (including soil crust, *S*) and bare, desert sites (*D*). (c) A two-dimensional spatial model can be constructed including these patch types, which are highlighted in (d). The local dynamics follows a set of stochastic rules that allow transitions between the three different states, as indicated in (e) using black arrows. Moreover, the presence of vegetation cover has an impact on these transitions, as indicated by the links ending in black circles. These are the basic rules used in previous literature modelling semiarid ecosystems [16, 17, 46]. The proposed scenario about synthetic terraformation of this system considered in this article is indicated using coloured links. The orange line denotes *C*-terraformation (a synthetic strain establishing cooperative feedbacks with vegetation), inhibiting the transition from *S* to *D*; the blue line indicates *D*-terraformation, allowing for increased dispersal of the engineered organisms (SYN) thus favouring the transition form *D* to *S*.

In a recent paper [47] it has been shown that, by tuning some particular features of models exhibiting catastrophic shifts, one can modify their nature and location in parameter spaces. Based on a theoretical approach, the authors suggested that the amount of stochasticity (both demographic and intrinsic) or population dispersal range can play that role. What type of microscopic mechanisms could actually avoid green-desert transitions? In this paper we seek using the insight gathered from models of dryland dynamics to predict the impact of bioengineering strategies on endangered drylands. In this context, several bioremediation approaches have been developed in the last two decades to enhance and stabilise soil biocrusts (for a overview, see [3] and references therein). These include a diverse range of approaches addressed to improve moisture and soil texture, from enrichment of key nutrients [22] or enrichment of cyanobacteria [28] to large-scale straw checkerboard barriers used to stabilise sand dunes [21].

More recently, ecosystem *Terraformation* has been suggested as a novel approach against green-desert shifts [24, 42, 43]. In a nutshell, some existing microorganisms, such as cyanobacteria from the soil crust, could be minimally modified by means of synthetic biology techniques to help improving soil moisture and create a cooperative feedback between vegetation or moss cover (see [13]) and the soil microbiome [27]. This engineering scenario aims at building *synthetic soil* e.g., soil crust, where the un-derlying community structure incorporates the synthetic strain. By doing so, potential tipping points could be made much more difficult to be achieved [44]. What might be the impact of these strategies on the large-scale, long-term dynamics of semiarid ecosystems? How the introduction or enhancement of ecological interactions (e.g., cooperation and microbes dispersal capacities) can modify the presence of tipping points? As shown below by means of both mathematical and computational models, the engineering of microorganisms present in arid and semiarid soils may easily expand the potential for ecosystems’ persistence.

## II. MODELLING TERRAFORMATION OF SEMIARID ECOSYSTEMS

In order to predict the impact of synthetic bioengineering strategies in arid and semiarid ecosystems, a spatial model given by a stochastic cellular automaton (CA) is introduced. Let us first consider the microscopic rules associated with the dynamics, as described by a set of transitions among the three defined ecological states. The basic transition diagram is displayed in Fig. 1(c), which includes the three potential states characterising a given patch, namely

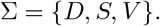

These states, as mentioned, are defined by desert (*D*) patches, by fertile soil (*S*) patches occupied by an engineered microorganism with their natural community (e.g., soil crust), and by a vegetated (*V*) state, respectively. This is of course an oversimplification that ignores most of the complexity and diversity involved, but allows for an analysis of the dynamics arising from the most fundamental ecological interactions. The model described here is an extension of the model proposed by Kéfi *et al.* (2007) [17] with an additional extra state associated to patches that are not occupied by plants but by soil crust plus (engineered) microorganisms. As discussed in a previous work [44], we aim at describing how the use of an appropriate synthetic design can help maintaining the stability and resilience of semiarid ecosystems.

The first rule is applied to degraded patches. Transition from degraded to fertile soil containing synthetic strains will take place with a probability:

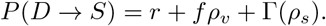

Here, parameter *r* is the rate of spontaneous restoration of fertile soil (due to, e.g., increased humidity, accumulation of organic matter, ecosystem’s engineers action, etc), and *f* is the influence of the surrounding vegetation in helping the transition to occur via a facilitation process. The constant *ρ*_*v*_ is the local density of vegetation. Finally, Γ(*ρ*_*s*_) denotes the spreading capacity of the microorganisms associated to their synthetic engineered properties (see below).

The second transition considers the reverse situation, namely a soil-to-desert transition:

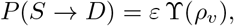

where *ε* is the rate of loss of fertile soil due to increased aridity or to other soil degradation processes (for simplicity we will study the range 0 ≤ *ε* ≤ 1). In this paper we consider (see below) a modification of the degradation rate mediated by a vegetation-dependent function ϒ(*ρ*_*v*_). The third rule involves the colonization of available *S* patches by vegetation:

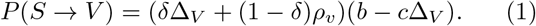

This term has the same form as the colonisation used in Ref. [17]. Here *δ* is related to seeds dispersal, and the last term in the r.h.s. of Eq. (1) stands for germination rate (balance between seeds that arrives to an unoccupied space that already have soil-crust (*S*), the germination rate and the degradation of seed). The parameter *δ* balances the influence of the local and global vegetation to produce the germination of new plants in a given site. Seeds are produced and germinate at rate *b*. However, they can also be degraded during the dispersal travel (at a rate *c*). Finally, a linear decay of vegetated patches to fertile soil occurs following a simple rule:

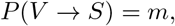

where *m* is the death rate of the plants, which can occur due to pathogens or the recollection by livestock activity.

Here two different but complementary terraformation strategies are considered and weighted by means of parameters *α* and *β*. Their role within the transition diagram is displayed in Fig. 1(c). They involve:

1. *C*-terraformation: Engineering cooperative loops between vegetation and soil organisms. Here a decreasing function of the vegetation cover, namely

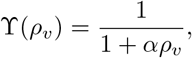

introduces a reduction in the impact of desertification due to increased soil quality favoured by the synthetic microbial population. The efficiency of this term is weighted by the constant *α*: large values of *α* imply a lower soil degradation rate due to the action of the synthetic microbes which are cooperatively coupled with vegetation at a local scale.
2. *D*-terraformation: Engineering the capacity of microbial spreading (e.g., faster replicative rates of the microbes, increased formation of endospores able to colonise local surroundings). The dispersal is here considered proportional to the term

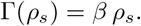

Here *ρ*_*s*_ is the local density of fertile soil. In other words, at a rate *β*, engineered strains improve their dispersal capabilities thus changing the properties of, i.e., desert patches by retaining humidity, depositing organic matter, etc.

Setting *α* and *β* to zero, the CA model recovers the original model introduced by Kéfi *et al.* (2007) [17].

## III. MODELS AND RESULTS

In the next sections we will explore the dynamics tied to the rules described above using both a well-mixed (mean field) and a discrete spatial setting using a stochastic cellular automaton (CA). The first approach, developed in Section A, employs differential equations to predict the presence of tipping points and changes in their location in the parameter space due to the effects of the addition of a synthetic strain. The second approach, studied in Section B, explicitly deals with a spatially extended population where local interactions take place on a two-dimensional lattice and the set of rules are applied in a probabilistic manner, thus considering stochastic effects. Finally, we provide insights into the nature of the tipping points found in the spatial model, focusing on the early warning signals for the system without and with synthetic engineering, a *Terraformed condition*.

### A. Mean-field model

The differential equations model, including the interactions and processes tied to the bioengineered synthetic strains, can be represented as follows:

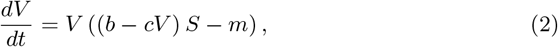

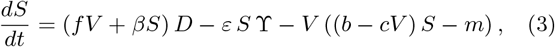

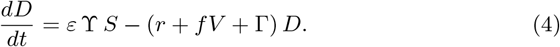

Equation (2) contains a logistic term that includes variable *S* as a multiplicative term, indicating that plants need viable soil to persist, and an exponential decay proportional to *m*. Equation (3) includes the positive effects triggered by vegetation and existing soil cover, as well as negative terms that are in fact symmetric to those present in the previous equation for *V*. Equation (4) provides the dynamics of desert patches resulting from desertification, recruitment to viable soil patch and an exponential decay from *S* to *D*.

Since the three classes of patches cover the entire lattice in the spatial model, it is possible to normalise and consider the states as population fractions in the mean field approach, i. e. *V*(*t*) + *S*(*t*) + *D*(*t*) = 1. This allows to reduce the three-variables system to a two-variables one using the linear relation

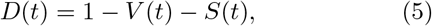

leading to:

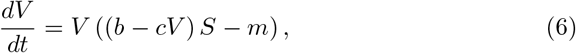

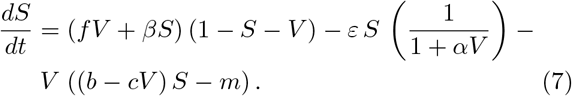

Under this model reduction, the fraction *D* is automatically obtained from Eq. (5) once the fractions of states *V* and *S* are determined. We must note that Eqs. (6)–(7) have been recently studied in Ref. [46] for the none engineered system (that is taking *α* = *β* = 0).

We are specially interested in those scenarios involving a full dominance of the desert state, focusing in the transitions between states. We have identified four equilibria for Eqs. (6) and (7), two of them implying the extinction of vegetation and another one allowing for the stable co-existence of the three ecological states. The first equilibrium is at the origin, i. e. 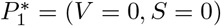, were the system becomes a desert (*D* = 1). The second equilibrium, keeps the fertile soil and the desert patches without vegetation, and is given by 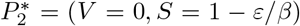. This state is interesting from the point of view of the terraformation strategies, since its existence depends on the chosen engineering strategy. If *β* = 0, when no engineered organisms are found in the soil crust, this equilibrium is not biologically meaningful (*S*(*t* → ∞) = −∞).

Two more equilibrium points, labeled 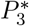 and 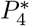, have been identified numerically (see Figs. S1-S4, and Figs.S6-223 S8 in the *Supplementary Information*). The equilibrium 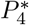 (numerical results suggest it is always stable within the simplex, see below) involves *V*(*t* → ∞) > 0. *S*(*t* → ∞) > 0 and *V*(*t* → ∞) + *S*(*t* → ∞) < 1, allowing the coexistence of the three ecological states when is an interior equilibrium. 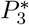 is a saddle point. See below and Figs. S1-S4 for further details on the dynamics of the equilibria on the simplex (*V,S*).

The conditions defining the local stability of each equilibrium can be obtained by means of the eigenvalues of the Jacobian matrix 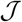, given, for Eqs. (6) and (7), by:

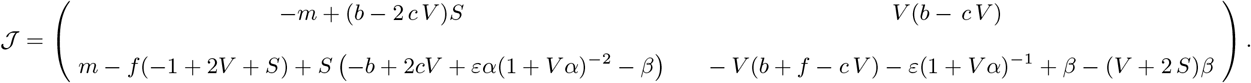

The eigenvalues of matrix 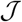 evaluated at an equilibrium provide its local stability conditions. The stability for equilibria 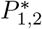 can be easily computed, while the stability of equilibria 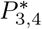 will be characterised numerically. In the case of the desert equilibrium 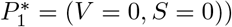, the eigenvalues are λ_1_ = −*m* and λ_2_ = *β* - *ε*. Since all the parameters are positive, this equilibrium will be stable when *ε* > *β*. For 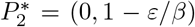, its eigenvalues are *λ*_1_ = *ε* - *β* and *λ*_2_ = − *m* + *b*(1 - *ε/β*), see Fig. S5 for a stability diagram for 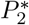 in the parameter space (*ε, β*). Note that 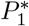 and 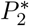 suffer a transcritical bifurcation at case of the desert equilibrium *P*1∗ = (*V* = 0*, S* = 0)), the *β* = *ε*, since the two conditions for this bifurcation are eigenvalues are *λ*1 = −*m* and *λ*2 = *β* − *ε*. Since all thefulfilled: the two fixed points collide at the bifurcation parameters are positive, this equilibrium will be stable value (at *ε* = *β*, 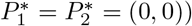, and they interchange the stability.

Figure 2 displays the critical degradation rate of the fertile soil, *ε*_*c*_, computed numerically in the parameter space (*α, β*), separating two domains with and without vegetation at equilibrium. The value of *ε*_*c*_ moves to higher values as either *α* or *β* are increased. The yellow region in the surface of *ε*_*c*_ in Fig. 2(A) denotes those values of *α* and *β* where *E_c_* ≥ 1 (although they are all set to *ε*_*c*_ = 1). This region denotes that no critical degradation is achieved under the chosen parameter values and under the restriction 0 ≤ *ε* ≤ 1. The four panels in Fig. 2(B) (corresponding to the values of *α* and *β* indicated in Fig. 2(A)) provide a more detailed picture of the changes in the fraction of the states due to both *C*- and *D*-terraformation strategies. Panel (a) shows how the states change without terraformation (*α* = *β* = 0) at increasing *ε*. Note that for *ε* > 0.2 the desert state becomes dominant. This critical degradation rate is displaced to larger values for the terraformed system, meaning that the ecosystem becomes much more resistant to soil degradation. More specifically, *ε*_*c*_ ≈ 0.3 when applying the *C*-terraformation strategy (panel (b) in Fig. 2(B)). This effect is further amplified for the *D*-terraformation, resulting in *ε*_c_ ≈ 0.7 (see panel (c) in Fig. 2(B)). The combination of both terraformation strategies (see Fig. 2(B) panel (d)) further increases the size of non-desert phases, here with *ε*_*c*_ ≈ 0.8. In both cases (specially for *D*-terraformation), the domain of fertile soil is increased due to the presence of the synthetic strains (Figs. S6, S7 and S8 show how the values of *ε*_*c*_ are displaced for different intensities of the proposed terraformation strategies).

**FIG. 2.**
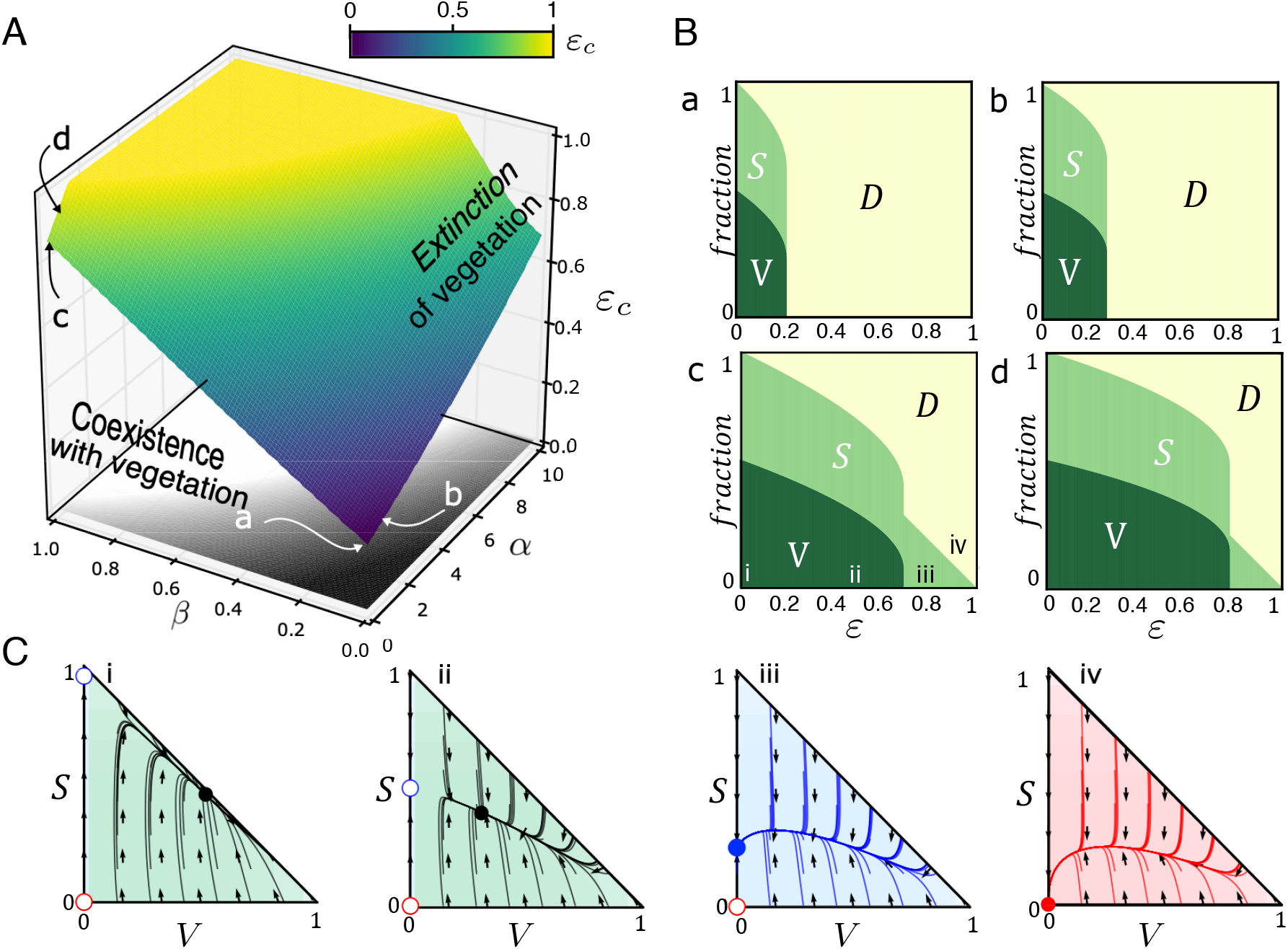
(A) Critical degradation rate of fertile soil (*εc*) computed in the parameter space of engineering strategies (*α, β*) from Eqs. (6)–(7). The inner surface separates the values of *ε* allowing for the presence of vegetation. The yellow area indicates those pairs of *α* and *β* giving *ϵ*_c_ ≥ 1 (those values of *ε*_c_ > 1 are set to *ε*_c_ = 1). (B) Fraction of the states at equilibrium increasing *ε*_c_ and using a full vegetated system as initial conditions *V* (0) = 1, *S*(0) = *D*(0) = 0. Panel (a) shows results for a non-engineered ecosystem (*α* = *β* = 0), and panel (b) for the engineered ecosystem incorporating cooperation loops (*α* = 1 and *β* = 0). Finally, panels (c) and (d) display, respectively, the results engineering the resilience of the soil crust (*α* = 0 and *β* = 1) and both strategies (*α* = *β* = 1). (C) Phase portraits for the case *α* = 0 and *β* = 1, with: (i) *ε* = 0.0; (ii) *ε* = 0.2; (iii) *ε* = 0.6; and (iv) *ε* = 0.9. The circles indicate the fixed points (stable: solid; unstable: open). We note that in the phase portraits (i) and (ii) there exists an interior saddle (see Figs. S1-S4 for the identification of the nullclines and dynamics). Green, blue, and red regions of the phase portraits denote the equilibrium states of *V* -*S* coexistence, *S*-*D* coexistence, and full desert, respectively. The other parameters are *r* = 0, *f* = 0.9, *δ* = 0.1, *b* = 0.6, *c* = 0.3, and *m* = 0.15.

The dynamics tied to increases of soil degradation rate are summarised in Fig. 2(C) (see also Figs. S1-S4), where different phase portraits are displayed for the values of *ε* indicated in 2(B) panel (c) (from (i) to (iv)). The transition from the coexistence of vegetation and fertile soil to no vegetation is catastrophic, and is due to a saddle-node bifurcation between fixed points 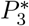 and 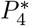 (transition between phase portraits (ii) and (iii), see also Figs. S1-S4). Once this bifurcation takes place, the only stable point is 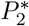 (with fertile soil and desert areas). Then, further increase of *ε* involves the transcritical bifurcation previously mentioned, after which the origin becomes globally stable and the desert becomes the only possible state (transitions between phase portraits (iii) and (iv) of Fig. 2(C)), see also Fig. S7 where the transcritical bifurcation is shown.

Finally, Fig. S6 provides one-dimensional bifurcation diagrams tied to the increase of *ε* for the non-terraformed system and the three possible terraformation strategies (*C*-, *D*-, and (*C, D*)- terraformation). Here we want to emphasise the presence of so-called delayed transitions (also called ghosts), which arise just after a saddle-node bifurcation takes place [20]. This is a dynamical phe-nomenon that involves extremely long transients once the bifurcations has occurred, and the time trajectories experience a long bottleneck before rapidly achieving, in Eqs. (6)–(7), another attractor (e.g., the full desert state in Fig. S6(a)-(b)). These transients are typically found in systems with strong feedbacks e.g., cooperation, such as catalytic hypercycles (see e.g., [32, 33]) or metapopulations with facilitation [41]. Also, this dynamical delay tied to saddle-node bifurcations has been recently described in both deterministic and stochastic well-mixed approaches for the non-terraformed system explored in this article (see [46] for more details).

### B. Spatial stochastic model

The mean field model studied in the previous section provides a first approximation to understand the qualitative dynamics arising from the nonlinear interactions of the studied system and the resulting tipping points, especially when including the proposed terraformation strategies. However, in order to test its robustness in a more ecologically-realistic setting, one has to take into account both local spatial correlations and stochastic effects. To do so, we build a spatially-explicit simulation model given by a stochastic cellular automaton (CA). The CA incorporates the previously studied states (*V*, *S*, *D*) and their transitions, which are now probabilistic. Spatial degrees of freedom are introduced by using 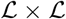 a square lattice with periodic boundary conditions. At each time step, the following four transition probabilities defining the rate of change for each site *k* of the lattice (see Fig. 1) are applied in an asynchronous manner:

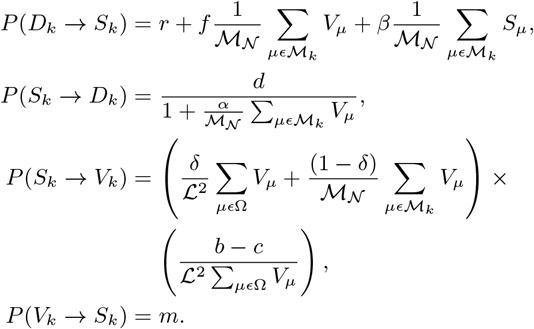

Here, 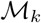 is the neighbourhood of site *k* and 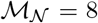 the number of neighbour (using a Moore neighborhood). The nature of local interactions is known to largely influence ecological dynamics. This is particularly important in drylands, where carbon and water limitation deeply constraints the outcome of nonlinear exchanges, leading to spatial patterning. Also, facilitation processes (involving strong nonlinearities) are known to introduce important changes in spatial systems, as compared to well-mixed ones. In this sense, recent research has found a shift from catastrophic tipping points to continuous ones due to local spatial processes [41, 47].

Similar analyses to those presented in Fig. 2 for the mean field model are displayed in Fig. 3 for the CA simulations. For the sake of comparison we have kept the same parameter values of the mean field model, implemented as probabilities in the CA, also using as initial conditions a full vegetated lattice. The critical surface separating the parameter scenarios allowing the persistence of vegetation are displayed in Fig. 3(A). Below the surface, *V* persists while above the surface it is only possible to find states *S* and *D*. Here, similarly to the mean field model, the yellow region indicates those values of *α* and *β* for which no critical degradation rate of the fertile soil is found (i.e., *ε*_*c*_ = 1). Note that the surface for the spatial system differs from the one obtained with the mean field model. This effect, taking into account that the lattice has 4 × 10^4^ sites (large system size), is probably introduced by space more than by stochasticity. This change of the surface is especially visible for parameter *α*. This may be explained because soil degradation rate (*ε*) makes the vegetation to decrease globally, but due to the local interactions of facilitation [48], plants remain in the ecosystem for larger rates of soil degradation.

**FIG. 3.**
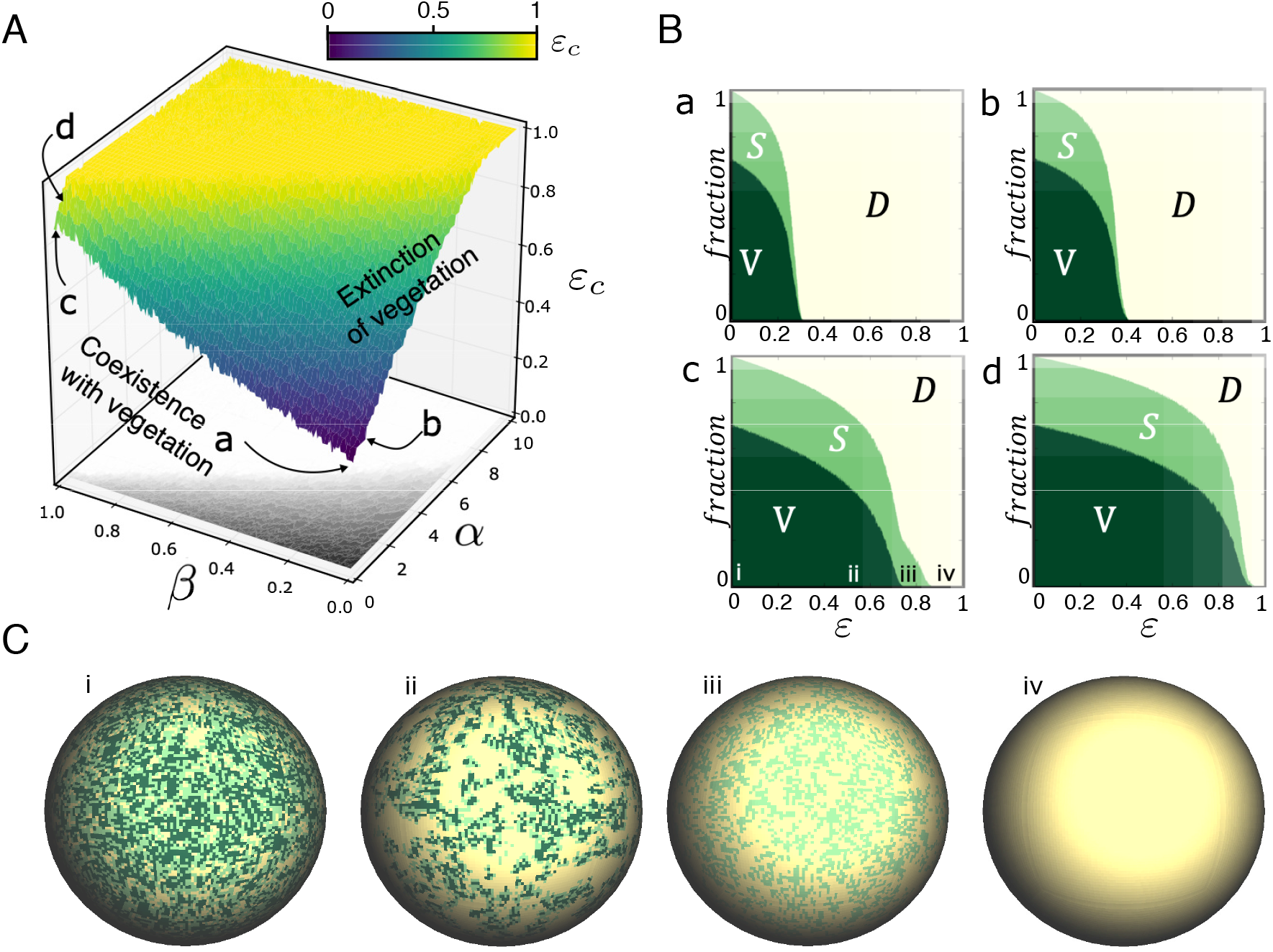
Same as in Fig. 2 for the spatial model. Vegetation extinction surface depending on the engineering strategies and the rate of fertile soil degradation (*ε*). The yellow region indicates that vegetation extinction can not be achieved because *ε* = 1. (B) Bifurcation diagrams depending on the engineering strategies (*α β*) and the degradation parameter *ε*. (a) Non-engineered ecosystem (*α* = *β* = 0), (b) engineering of cooperative loops between synthetic microorganisms and the vegetation (*α* = 1 and *β* = 0), (c) engineering resilience of the soil crust (*α* = 0 and *β* = 1); and (d) engineering both terraformation strategies (*α* = *β* = 1). The other probabilities are fixed using the same values of the parameters from Fig. 3. In all of the analyses the initial condition used is *V* (0) = 1. All these results have been obtained using a lattice with 200 *×* 200 sites.

The fraction of the three states computed at increasing *ε* is displayed in Fig. 3(B). All diagrams (a-d) display the same phenomenon: the introduction of spatial correlations shifts the extinction of the vegetation to larger values of *ε*. Consistently with mean field results, both the *C*- and *D*-terraformation displace the value of *ε*_*c*_ to larger values, being the (*C, D*)-terraformation strategy much more efficient in doing so under the selected parameters. The spheres displayed in Fig. 3(C) show the spatial patterns at equilibrium for the regions labeled in-panel (c) of Fig. 3(B), with: (i-ii) coexistence of the three states; (iii) only fertile soil and desert patches; and (iv) the full desert scenario.

In order to identify the nature of the transitions tied to the increase of *ε*, we have computed the stationary values of the fraction of areas with vegetation and fertile soil. We recall that both mean field and well-mixed stochastic simulations for the system without terraformation revealed a catastrophic transition and delayed transitions due to a ghost [46]. Also, the mean field model studied in Section III A have revealed the presence of saddle-node bifurcations responsible of vegetation extinctions. Figure S9 shows these results together with the spatial patterns on the square lattice and time series. Specifically, Fig. S9(a) shows results increasing *ε* for the system without terraformation. For this case *ε*_*c*_ ≈ 0.24. The same results are displayed by setting *α* = *β* = 1, for which the critical degradation moves to *ε*_*c*_ ≈ 0.85. For both cases, the discontinuity of the transition is not so evident as in the mean field model, even looking like a continuous one.

To identify the transition governing vegetation extinction we have used an indirect method analysing the properties of the dynamics close to the transition value. The results displayed in Fig. S10 indicate the presence of bottlenecking phenomena tied to the extinction of the vegetation, meaning that the transition is governed by a saddle-node bifurcation. We must notice that we have used a lattice of side size 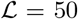 for these analyses due to the huge computational cost of computing extinction transients in large spatial systems. Figure S10(a) shows the transition for the non-engineered system. Specifically, panel (a.1) displays 5 realisations obtained with *ε* = 0.26, and the bottleneck region can be clearly seen (see Fig. S6 for comparison with the mean field bottlenecks). The delaying effect of the ghost can also be seen in panels (a.2), (a.3) and (a.4), which indicate the values at which vegetation spends longer iterates before achieving the full desert state. The same results are shown for the system with (*C, D*)- terraformation. Here, the extinction dynamics of the vegetation (investigated setting *ε* = 0.86) is also given by a clear bottleneck. Previous research in catalytic hypercycles also revealed extinctions due to saddle-node bifurcations and the corresponding bottlenecking phenomena in a CA model [33].

As discussed above, the approach to tipping points is often associated to a qualitative change in the nature of. the fluctuations exhibited by the system. How will our model behave when dealing with an extra phase associ-ated to vegetation-free, synthetic soil crust? A divergence in a stochastic dynamical system can be detected by using standard deviation measures. We have used several measures that can detect the approach to criticality in both the non-manipulated and the terraformed scenarios (spatial variance and distribution of patch sizes). The spatial variance *σ*_*ψ*_, with *ψ* referring to either *V* or *S*, is determined from

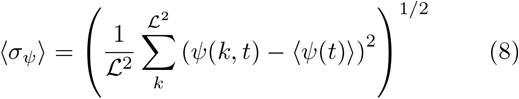

where we have used the mean over the entire lattice, given by

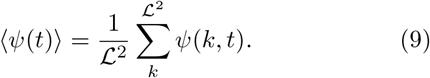

The fluctuations diverge in a characteristic fashion (Fig. (4)(a,b)): whereas *σ_V_* slowly grows and accelerates close to criticality, the same measure for *σ_S_* displays a slight decline only growing very quickly close to *ε*_*c*_. Despite these trends, both variables exhibit high fluctuations close to the transition to the desert state.

**FIG. 4.**
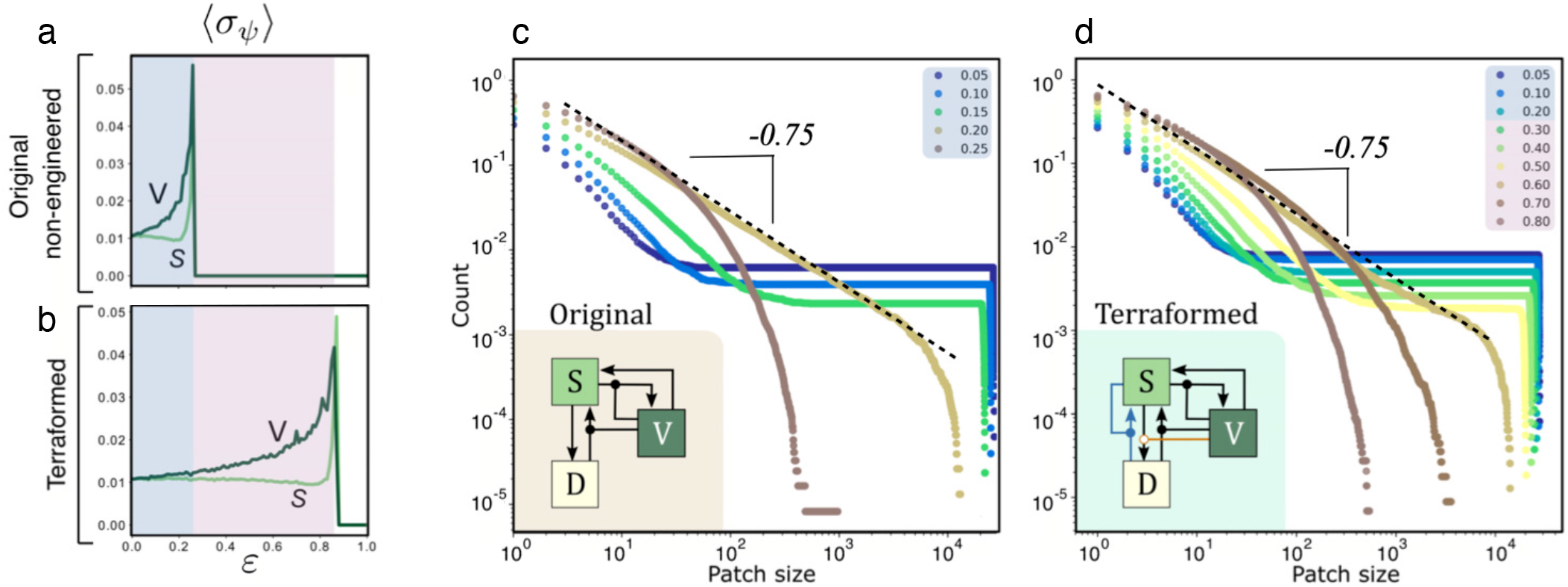
Warning signals for the non-engineered (*α* = *β* = 0) and terraformed (*α* = 1 and *β* = 1) ecosystems taking vegetation and fertile soil as indicators at increasing *ε*. Panels (a-b) display the standard deviation 〈*σ*_*ψ*_=*V,S*〉 of the population fluctuations for the original and the terraformation scenarios, corresponding to the simulations shown in Fig. S9. Values of *σ*_*ψ*_ have been obtained from the last 10^3^ values from time series with 10^6^ iterations. In (c-d) the cumulative distributions of patch sizes are displayed for different values of *ε* (indicated by the different colours inside the plots). The patch size distributions have been computed using 100 replicas from a lattice with 200 200 sites.

An additional confirmation of the criticality associated to the transition to the desert state is obtained by determining the distribution of vegetation cluster sizes. In a nutshell, criticality is known to be linked to power law distributions of connected clusters. On particular instance of these scaling behavior was found in [17]. Clusters of size *S* are composed by a set of *S* connected sites (i. e. sharing the same state and all elements in contact with another as nearest neighbors).

The number of clusters of a given size and their distributions have been obtained using a burning algorithm [12]. If *P* (*S*) indicates the probability of finding a cluster of size *S*, most measured patch size distributions found in arid and semiarid habitats follow a general form described as a truncated power law, namely *P*(*S*) ∼ *S*^−*τ*^ exp(−*S/S*_c_). This defines a power-law decay with an exponent *τ* that characterises the class of phenomenon underlying the dynamics [25] and an upper limit introduced by a cut-off *S_c_* in the exponential decay term. Close to critical transitions we should expect 462 a fat-tailed decay in *P* (*S*) following the power law form *P*(*S*) ∼ *S*^−*τ*^ (when no characteristic scale is present, i. e. for *S*_*c*_ → ∞). Instead, far below the transition point, the exponential term dominates and a characteristic scale is observed. In order to improve the smoothness of the distribution, the so called *cumulative* distribution *P*_>_(*S*) will be used, and is given by

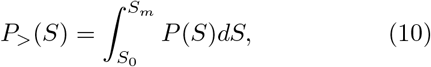

where *S*_0_ and *S*_*m*_ indicate the smallest (the one-site scale) and largest (lattice) sizes. At criticality, the cumulative form scales as a power law

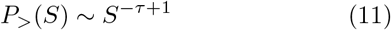

In our model, the shapes of the distributions change at increasing values of *ε*: from single-scaled to scale-free, as shown in Fig. 4(c) (original model) and Fig. 4(d) (terraformed). In both cases the same pattern if found, but the power-law behavior is observed at much higher levels of soil degradation. Interestingly, the scaling exponent for both cases is the same: *τ* ≈ 1.75, in agreement with the results reported in [16]. The consideration of the extra processes tied to terraformation do not alter a universal behaviour pattern, thus suggesting that, despite the parameter shift in *ε*_*c*_, the warning signals associated to both scenarios will behave in a very similar way.

## IV. DISCUSSION

The impact of anthropogenic-driven processes is rapidly increasing, leading to a degradation of extant ecosystems and in many cases pushing them closer to viability thresholds [1, 2, 14, 34]. In the case of semiarid ecosystems, this degradation is produced mostly by global warming and intensive grazing [15]. The increasing pressure could end up in catastrophic transitions that are essentially irreversible. It is expected that the expansion of arid areas will increase in the next decades, with an associated increase in the likelihood of green-desert tipping points. Under this potential scenario, it is important to develop strategies of intervention aimed at avoiding these transitions.

Theoretical results on the nature and location of these shifts suggest that some generic factors (such as noise and dispersal) [41, 47] could be crucial. Can realistic interventions help preventing green-desert transitions? As discussed above, several bioremediation strategies to face this problem have been proposed. Some recent proposals suggest how to exploit nonlinear features of tipping points based on periodic replanting [46]. More recently, an engineering approach based on synthetic biology has also been suggested [42–44]. In this article we investigate this latest approach for arid and semiarid ecosystems employing both mathematical and computational models.

Beyond the standard vegetated-desert model, in our study, we pay attention to the possibility that an engineered soil crust (a “synthetic” soil) might have on the phase space of potential ecosystem-level states. The original motivation is to describe, using a toy model, the impact two different but complementary strategies: the so-called *C*-terraformation using a designed cooperative loop (promoted by a designed microbial strain); and the *D*-terraformation, consisting on engineered strains with increased dispersal capacities. We are especially interested in the impact of these two synthetic terraformation strategies on the location and behaviour of tipping points.

Both terraformation strategies are shown to increase the resistance to soil degradation, pushing the tipping points to higher (or even much higher) critical degradation rates (*ε*_*c*_). This is consistently predicted by both mean-field and spatial models. However, when space is explicitly included, the parameter space supporting this protection against catastrophic shifts is much larger, with a fluctuation dynamics (and underlying warning signals) that behave similar between the non-engineered and the engineered systems i.e., they show the same universality pattern.

Our theoretical predictions are limited by the simplifying assumptions that define our model. In particular, the description of soils ignores their development, spatial organisation or diversity. These are multi-species communities, describable as complex networks [26]. These webs are known to display their own critical thresholds and thus it remains open how such diverse communities may impact the results reported here. However, despite all these limitations, it is important to highlight that previous models of vegetation dynamics in drylands have been very successful in providing deep insight into their natural counterparts [18]. Some key universal properties might pervade the success of these models. In our context, this is likely to be related to the general patterns found close to phase transitions. Additionally, the terraformation scheme described here is likely to be effective when the cooperative interaction helps creating the proper ecosystem engineering effect. By engineering ecosystem engineers, endangered arid and semiarid habitats might get protected (at least temporarily) from sudden collapse.

## Ethics

N/A.

## Data access

This article has no additional data.

## Author contributions

All authors built and analyzed the mathematical models. All authors wrote the paper and gave final approval for publication.

## Acknowledgements

The authors specially thank Sergi Valverde for the spatial rendering software with the cellular automata. We also thank Nuria Conde, Fernando Maestre and Miguel Berdugo for helpful discussions.

